# Contribution of the collective excitations to the coupled proton and energy transport along mitochondrial crista membrane in oxidative phosphorylation system

**DOI:** 10.1101/2022.11.16.516755

**Authors:** Semen V. Nesterov, Lev S. Yaguzhinsky, Raif G. Vasilov, Vasiliy N. Kadantsev, Alexey N. Goltsov

## Abstract

The results of many experimental and theoretical works indicate that after transport of protons across the mitochondrial inner membrane (MIM) in oxidative phosphorylation system (OXPHOS), they are retained on the membrane-water interface in non-equilibrium state with free energy excess due to low proton surface-to-bulk release. This well-established phenomenon suggests that proton trapping on the membrane interface ensures vectorial lateral transport of protons from proton pumps to ATP synthases (proton acceptors). Despite the key role of the proton transport in bioenergetics, the molecular mechanism of proton transfer in the OXPHOS system is not yet completely established. Here, we developed a dynamics model of long-range transport of energized protons along the MIM accompanied by collective excitation of localized wave proragating on the membrane surface. Our model is based on the new data on the macromolecular organization of OXPHOS showing the well-ordered structure of respirasomes and ATP synthases on the cristae membrane folds. We developed a two-component dynamics model of the proton transport considering two coupled subsystems: the ordered hydrogen bond (HB) chain of water molecules and lipid headgroups of MIM. We analytically obtained two-component soliton solution in this model, which describes the motion of the proton kink, corresponding to successive proton hops in the HB chain, and coherent motion of a compression soliton in the chain of lipid headgroups. The local deformation in a soliton range facilitates proton jumps due to water molecules approaching each other in the HB chain. We suggested that the proton-conducting structures formed along the cristae membrane surface promote direct lateral proton transfer in the OXPHOS system. Collective excitations at the water-membrane interface in a form of two-component soliton ensure the coupled non-dissipative transport of charge carriers and elastic energy of MIM deformation to ATP synthases that may be utilized in ATP synthesis providing maximal efficiency in mitochondrial bioenergetics.

## 1. Introduction

One of the main bioenergetics issues is still not fully resolved – how the efficient energy transfer in mitochondrial oxidative phosphorylation (OXPHOS) system between proton pumps and ATP-synthases is achieved. While Mitchell’s chemiosmotic theory provides general understanding of the OXPHOS system functioning, it ignores a large number of experimental facts showing that after transmembrane transfer or after H^+^ dissociation at the membrane-water interface, protons do not immediately equilibrate with a water bulk phase, but are retained at the membrane-water interface in non-equilibrium state (review in (Mulkidjanian et al. 2006; Nesterov et al. 2022)). The retention of the protons on the membrane was shown in the model bilayer membranes using different techniques for proton release (Antonenko et al. 1993; Tashkin et al. 2019). On the liposomes (Sjöholm et al. 2017) and even on the octane-water interface (Yaguzhinsky et al. 1976), it was shown that surface protons drive ATP synthesis. The origins of proton affinity for the interfaces were studied both experimentally (Weichselbaum et al. 2017) and theoretically (Lee 2021; Medvedev and Stuchebrukhov 2012; Cherepanov et al. 2004). The retarded membrane to bulk transfer of protons was shown in photosynthetic systems (Junge and Ausländer 1974), rhodopsin purple membranes (Drachev et al. 1984; Heberle et al. 1994) and mitochondrial inner membrane (Moiseeva et al. 2011). Not long ago it was experimentally shown that under operation of mitochondrial proton pumps there is no significant acidification of the intermembrane space (Toth et al. 2020) and there are also indications that there is a lateral proton concentration gradient between proton pumps and ATP synthases (Rieger et al. 2014).

Despite all of the above, the full theory of proton transport on the interfaces is not yet concluded, so there is still a need to develop a satisfactory physicochemical explanation of the proton transfer mechanism in the OXPHOS system on the membrane surface taking into account local environment. It remains unclear how exactly proton-membrane interaction occurs, how much energy can be transferred by non-equilibrium protons and in what form, and how this energy is used by ATP synthase. In this work, the particular attention was given to the question of how fast proton motion along the membrane is ensured while the proton strongly interacts with membrane, even changing the structure of the membrane itself (Deplazes et al. 2018). At first glance, it should create high energy barriers for proton transfer or lead to significant energy dissipation. Indeed, in an artificial model system, when the distance between proton pump and ATP-synthase becomes more than 80 nm, energy dissipation led to the impossibility of ATP synthesis (Sjöholm et al. 2017).

One of the molecular mechanisms underlying the effective proton transport over long distance was considered to be Grotthuss-like mechanism of proton transfer (structure diffusion) in the hydrogen bond (HB) chains formed by water molecules (Marx 2006; Wraight 2006). This mechanism was applied to the description of proton conductivity in various molecular systems, i.e. transmembrane channels of bioenergetics proteins in which proton hops from one water molecule or titratable group to the next ones (DeCoursey 2003; Sakashita et al. 2020; Borshchevskiy et al. 2022). It is reasonable to apply the same approach to the consideration of the direct motion of a proton along the membrane, taking into account the latest experimental data that the proton motion occurs at the interphase along bound water (Weichselbaum et al. 2017), but not along the titratable groups (Springer et al. 2011). However, it is not excluded that membrane groups can modulate the water HB network formation.

It was recognised that the interaction of the ordered HB chains with their environment (amino groups in surface of protein channels and lipid polar groups at the membrane interface) plays a significant role in maintaining effectiveness of proton transfer (Mulkidjanian et al. 2006; Weichselbaum et al. 2017). The proton transport along the proton-conducting HB structures requires excitation of collective modes in these structures that cause in part self-assembling of water molecules into the ordered HB chains and its collective reorientation as a result of structural diffusion (hop-and-turn mechanism). Theoretical consideration of the proton transport as a collective phenomenon emerging in molecular systems has been first carried out in the framework of Davydov soliton theory (Antonchenko et al. 1983). Davydov soliton (Davydov 1982) or other similar self-localized states (Austin et al. 2003; Pang 2012) and collective excitations (Bolterauer et al. 1991; Kadantsev and Goltsov 2020) have long been and still are considered as promising candidates for the role of charge and energy carriers within proteins (Kavitha et al. 2016), along the lipid membrane surface (Manousakis 2005; Kadantsev and Goltsov 2022), and on interfaces of biopolymers and artificial membranes (Matsui and Matsuo 2020). The concept of solitary waves was developed to described the mechanism underlying highly effective transport of charge and energy in molecular systems in the form of collective excitations which are described by autolocalized solutions (solitons) of nonlinear wave equations (Davydov 1985). It was shown that solitons can propagate in quasi-one-dimensional molecular systems over long distance conserving their shape and energy (Davydov 1977). The further progress in this direction was made within different modifications of two-component soliton models, where interaction of proton motion with the hydroxyl ion vibration and dynamics of orientation defects in the HB chain were taken into consideration (Savin and Zolotaryuk 1992; Zolotaryuk et al. 2000; Lupichev et al. 2015). Note all these models were developed for the ice-like structures of the HB chains, and their application to biological structures requires further development and detailed consideration of the real structure of the proton environment effecting the proton transport. The comprehensive model of the soliton transport was developed and applied to the proton transfer along artificial assembled monolayer surface made up of sulfonic acid headgroups arranged in a regular hexagonal array (Golovnev and Eikerling 2012). In this model, relationship between the mobility of soliton and macroscopic parameters of the sulfonic acid array structure were established. The next step in this direction should be consideration of dynamic behaviour of the proton environment and its role in effectiveness of proton transport in biological interface structures.

This paper aims at the elucidation of the molecular mechanism underlying collective dynamics in proton transfer along the HB chains formed by ordered water molecules interacting with the environment - mitochondrial inner membrane (MIM). We based on the new structural data on the OXPHOS system obtained by cryo-electron tomography (Nesterov et al. 2021a), which allowed constructing a model reflecting structural, function, and physicochemical characteristics of the OXPHOS system and MIM. We suggested that the proton-conducting structures along the cristae membrane connect proton pumps (proton sources) with ATP synthases (proton sinks) and promote direct proton transfer in the OXPHOS system. We developed a dynamics model of long-range transport of energized proton along MIM, which is accompanied by collective excitation of localized waves propagating along the membrane and facilitating proton transfer. Coherent motion of proton in the HB chain and local deformation wave in MIM was described in a two-component soliton model. According to the developed model, the two-component soliton ensures coupled dissipationless transport of charge and elastic energy of MIM deformation to ATP synthases that may be utilized in ATP synthesis and control mitochondria bioenergetics.

## 2. Model background and formulation

### 2.1. The OXPHOS system organization in mitochondrial cristae membrane folds

Development of the dynamics model of proton transport in the OXPHOS system should reflect the real structure of that system. In our model, we used the new data on the macromolecular organization of the OXPHOS system and complex cristae ultra-structure obtained by cryo-electron tomography (cryo-ET) (Davies et al. 2011; Nesterov et al. 2021a) (Fig. 1A). The most ordered known forms of the OXPHOS system are located on the cristae folds and consist of clustered oligomeric structures of parallel rows of respirasomes and rows of ATP synthase dimers. Even in less compressed and ordered structures, ATP synthase dimers form rows at the edges of the cristae (Strauss et al. 2008). This makes possible to consider proton transport as unidirectional motion from the pump row to the ATP synthase row (Fig. 1B). The short distance of less than 80 nm between the observed rows of proton pumps and ATP synthase dimers provides the direct and fast transfer of proton to ATP synthases along the cristae membrane avoiding proton surface-to-bulk release (Sjöholm et al. 2017). The fact that hydrogen ions move laterally in a thin near-membrane layer of water (Weichselbaum et al. 2017) is also taken into account in our model. Schematic representation of proton transfer along the membrane from the respirasome to ATP synthase is shown on Fig. 1C. It should be mentioned that apart from charge carrier transport, pumping protons can also transfer the excess free energy due to their nonequilibrium hydration shell formed on the membrane-water interface (Nesterov et al. 2022). The excess free energy of the energized proton may cause local membrane deformation and induce collective excitation formation, the energy of which along with the proton is then utilized by ATP synthase.

**Fig. 1.**
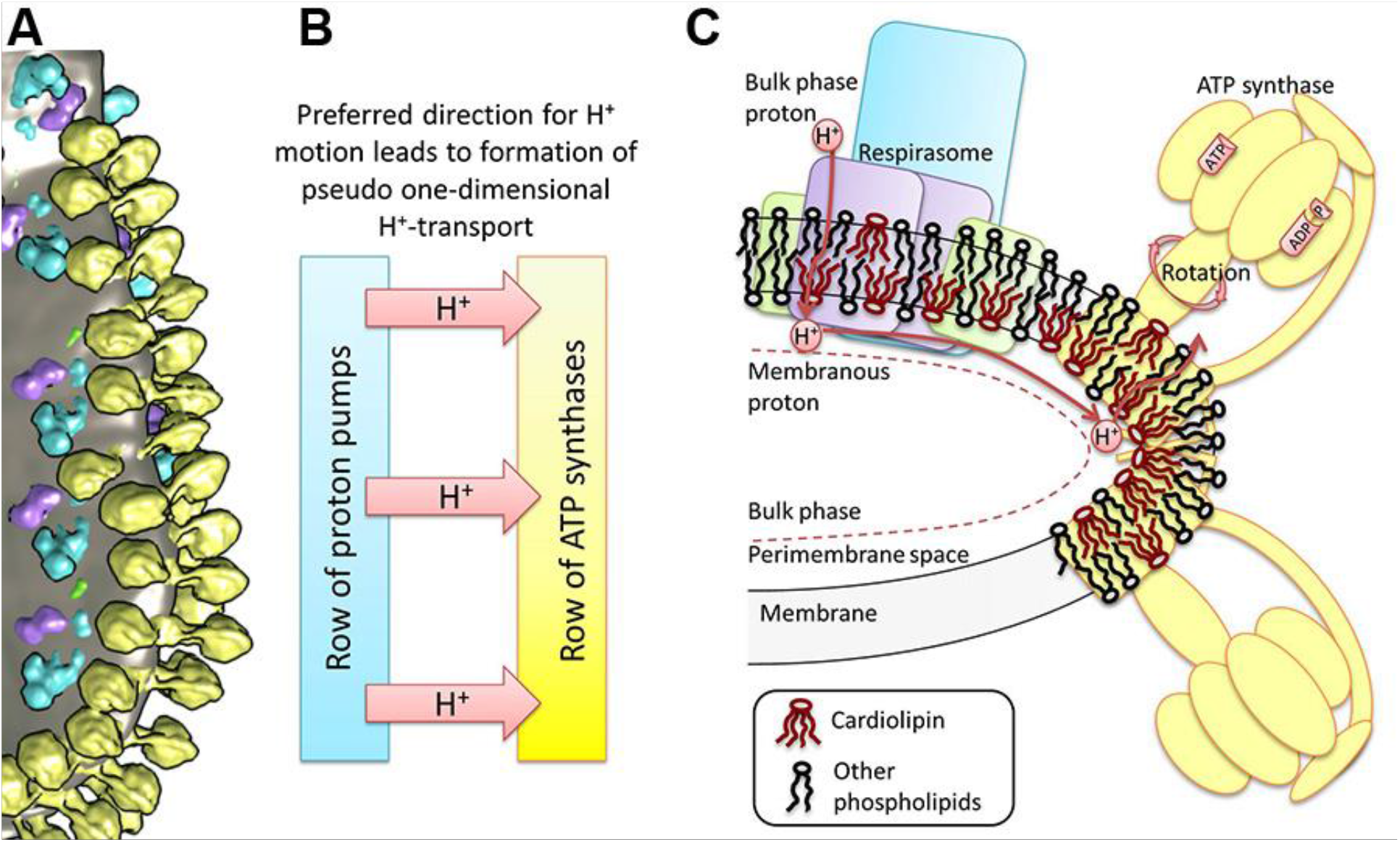
Structure of mitochondrial OXPHOS system and cristae membrane illustrating a proton transfer pathway. (A) The cluster of components of the OXPHOS system at the bend of crista of heart mitochondria (Nesterov et al. 2021a). Yellow – ATP synthase dimers, blue – complex I, purple – complex III dimers, green – complex IV, grey – lipid membrane. (B) A dedicated direction of proton transfer between rows of proton pumps and ATP synthases. (C) Schematic reconstruction of the cluster in the OXPHOS system on the membrane fold and a pathway of the lateral transfer of protons from the respirasome to ATP synthase. The area of increased curvature of the membrane is enriched with CL molecules.

### 2.2. Role of cardiolipin in cristae membrane structure, dynamics and function

The folds of the cristae are enriched with a lipid specific for the MIM – cardiolipin (CL), which, due to its conical shape, accumulates in areas with a high curvature of the membrane (Beltrán-Heredia et al. 2019). CL molecules selectively interact with respiratory chain complexes (Arias-Cartin et al. 2012; Arnarez et al. 2013), mitochondrial transporters (Beyer and Klingenberg 1985) and ATP synthases (Gasanov et al. 2018; Mühleip et al. 2019). Moreover, CL not only binds to components of the OXPHOS systems, but also is essential for their functioning. When CL in submitochondrial particles is damaged by reactive oxygen species (ROS), complexes I, III, and IV become dysfunctional, while the addition of CL restores their functions (Paradies et al. 2002, 2001, 2000). CL is also involved in the functioning of ATP synthase (Gasanov et al. 2018; Garab et al. 2022), nucleotide translocator (Kunji and Ruprecht 2020), and is also necessary for the assembly and functioning of respirasomes (Zhang et al. 2002, 2005; Pfeiffer et al. 2003). Thus, the areas of high curvature of MIM, where the clusters of the OXPHOS system are located, are enriched in CL which is involved in the operation of nearly all OXPHOS system components (Joubert and Puff 2021).

CL possesses at least one or two strong acid phosphate groups (Kooijman et al. 2017), capable of hydrogen bond formation with water or hydronium molecules. Due to CL importance for the OXPHOS system and its ability to influence water orientation in the near-membrane zone, we considered its presence and special properties when constructing the model of proton transfer in the OXPHOS system. It is known that CL does not form covalent bonds with positively charged lysine residues of ATP synthase rotor, although it selectively interacts with them (Duncan et al. 2016). Thus, CL acts as a lubricant, preventing the ATP synthase rotor from strong chemical bonds formation with the environment. It is possible that CL can perform a similar role to ensure the motion of positively charged hydronium.

### 2.3. Grotthuss-like mechanism of proton transfer along the HB chains

As was mentioned in the Introduction, one of the molecular mechanisms underlying the effective proton conductivity is Grotthuss-like mechanism of proton transfer in the HB chains which is applicable both to transmembrane channels and water-membrane interface systems. In our model, we considered Grotthuss-like mechanism of proton motion in the HB chains formed by water molecules strongly hydrogen bonding with the phosphate oxygens at the water-lipid interface (Fig. 2).

**Fig. 2.**
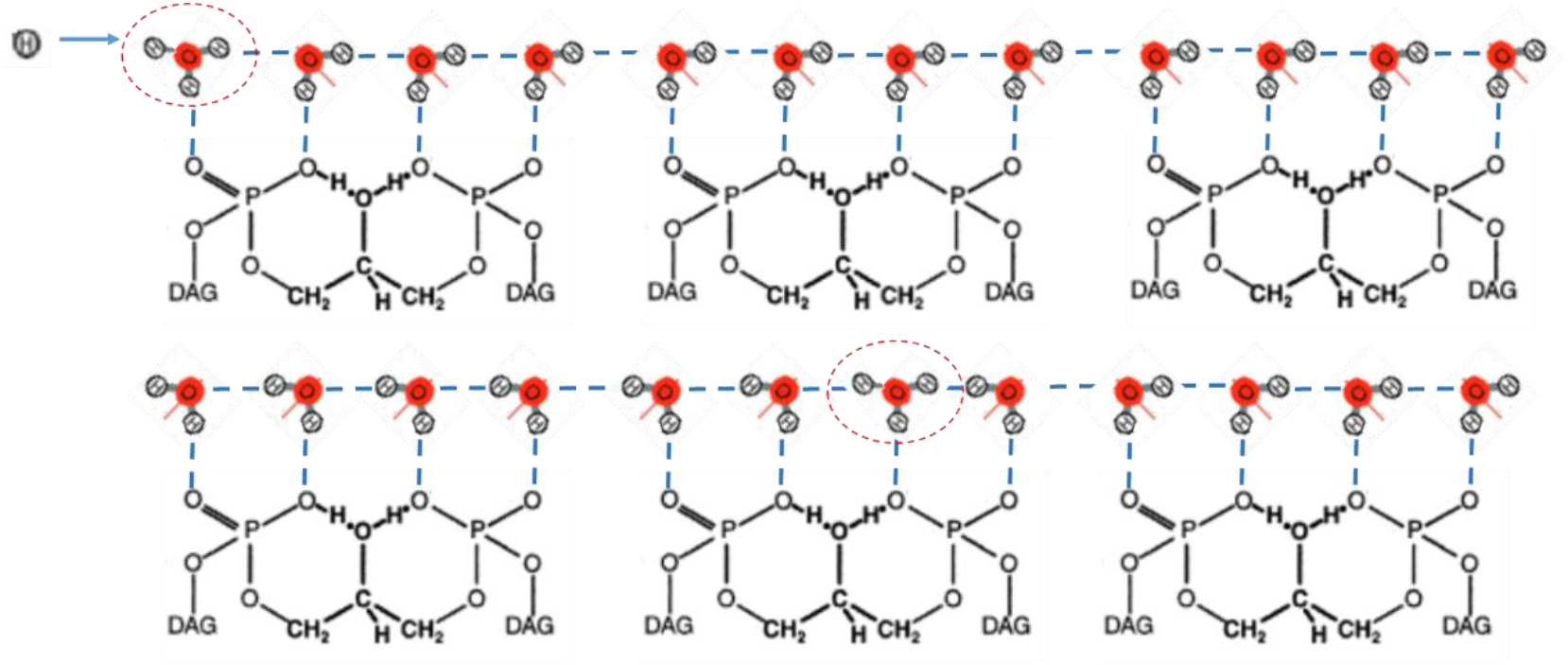
Proton transport along Grotthuss wires formed by the HB chain of the oriented water molecules hydrogen bonding with the phosphate oxygens of CL polar headgroups. Upper scheme shows an initial orientation of the HB chain with hydronium (encircled) in the beginning of the chain; bottom scheme shows propagation of proton (hydronium) and a change in water molecules orientation. Blue dash lines - hydrogen bonds. Red line – orientation of the dipole moment of water molecules. DAG – diacylglycerol group of CL.

As was mentioned above, the MIM and especially the vicinity of the OXPHOS system are enriched with CL molecules. We proposed that the HB chain of water molecules may be assembled on the surface of CL domains which were observed in MIM (Mileykovskaya and Dowhan 2009), model membranes (Beltrán-Heredia et al. 2019), and were investigated by computational modelling in curved two-component membranes (Huang et al. 2006). While proton transfer occurs along the water HB chains, the positioning of water molecules and distances between them are controlled by water-lipid interactions. Therefore, water-lipid interaction on the interface makes it possible to consider the connection between proton motion and elastic deformation of the membrane.

### 2.3. A two-component model of lateral proton transport along mitochondria crista membrane

In this section, we propose a molecular mechanism of long-range proton transport along the mitochondrial inner membrane, which is facilitated by the acoustic soliton moving along the membrane. Assuming molecular structure of the MIM which was described above, we developed a two-component dynamics model of the proton transport. The model includes two subsystems. The first one is the HB chain of water molecules bound to the second subsystem including the CL headgroup chains within CL-enriched domains. For the first subsystem, we wrote the Hamiltonian of the proton in the HB chain in the following form

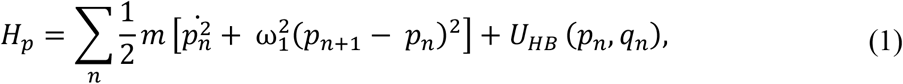

where *p*_*n*_ characterizes the displacement of the *n*th proton relative to the middle of the *n*th HB. The point above *p*_*n*_ stands for differentiation by time *t*. 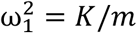, where *K* is the stiffness constant of the proton-proton interaction in the HB chain. Consideration of the interaction between adjacent protons provides cooperative behaviour and ordered states of the HB chain when protons are occupied either left or right well of the double-well potential *U*_*HB*_ (*p*_*n*_, *q*_*n*_). This proton-proton interaction defines two degenerate ground states, which are shown in Fig. 2.

The HB potential *U*_*HB*_ (*p*, *q*) is a two-dimensional potential surface depending on two variables, *p* and *q*, where *q* is the displacement of hydroxyl ions ОН^−^ from its equilibrium. It describes the energy potential of the proton, which is formed by the two neighbour hydroxyl ions OH^−^. A shape of the potential *U*_*HB*_ (*p*, *q*) and its dependence on *q* are defined by a distance between two neighbour hydroxyl ions, R. At distance *R* in a range of 2.5 – 2.6 Å, the potential *U*_*HB*_ (*p*, *q*) is double-well which is transformed into a single well potential when *R* is reduced below the critical value *R_c_*=2.5 Å (Ishikita and Saito 2014). In our model, we used the following approximation of the potential *U*_*HB*_ (*p*, *q*)

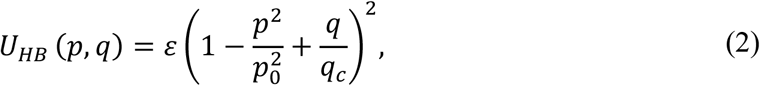

where potential parameters *ε*, *p*_0_ and *q*_*c*_ define its shape (Antonchenko et al. 1983; Zolotaryuk et al. 2000). At *q* > −*q*_*c*_, the potential is double well and represents two proton degenerative equilibrium positions at 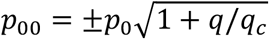 separated by the potential barrier of height *ε*. At ≤ −*q*_*c*_, the double-well potential is transformed to a single well potential. This transformation occurs when the distance between two hydroxyl ions is reduced to less than critical displacement *q*_*c*_. Parameters of *U*_*HB*_ (*p*, *q*) (2) were determined based on the approximation of the quantum chemistry calculation of the HB potential by the φ^4^-function for the hydrogen bonding hydronium and water molecules at various distances between them (Duan et al. 1993). The HB potential with parameters *ε* = 10.5 kJ/mol, *p*_0_ = 0.26 Å and *q*_*c*_ = 0.2 Å describes well a potential change when distance between hydronium and water molecules changes from 2.3 Å to 3 Å. The HB potential surface *U*_*HB*_ (*p*, *q*) (2) at the defined parameters is shown in Fig. 3.

**Fig. 3.**
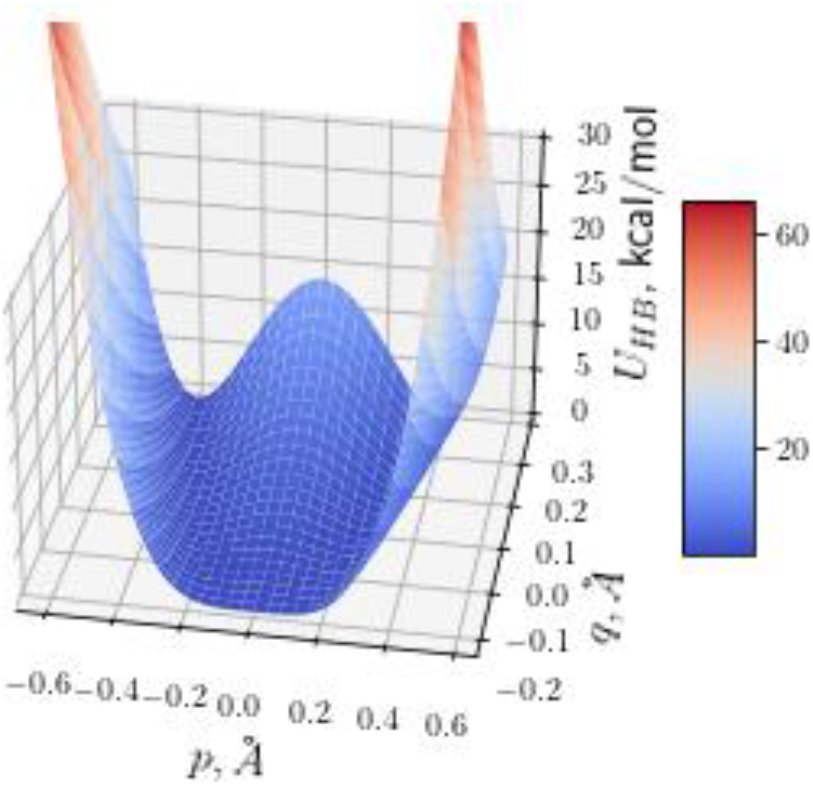
The HB potential *U*_*HB*_ (*p*, *q*) depending on the positions of proton *p* and OH^−^ ions *q*.

In the model, we considered Grotthuss-type conduction mechanism of proton in the HB chain using the double well potential (2). Proton transfer is a fast exchange of proton between hydronium (donor) and water molecules (acceptor) that correspond to successive proton jumps from one equilibrium position to another generating hydroxyl and hydronium ions (Fig. 2). Proton jumps occur through potential barrier, which depends on the displacement *q* between neighbour OH^−^ ions according to Eq. (2). The effective height of the barrier decreases with decreasing displacement *q* as

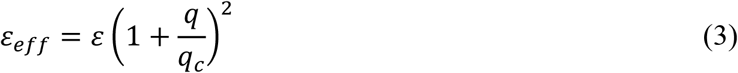

at *q* < 0. The lowering the potential barrier facilitates proton transfer in a local region of compression in the HB chain. To consider the motion of OH^−^ ions, we included in the model dynamic description of the CL molecules.

The Hamiltonian of interacting CL in the harmonic approximation may be written as follows:

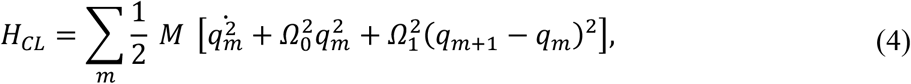

where *M* is CL mass; Ω_0_ and Ω_1_ are the characteristic frequencies of vibrations in the lipid system. The last term in Eq. (4) takes into account dispersion of elastic waves in the CL chain.

The Hamiltonian of the whole “HB chain + CL chain” system may be written in the continuum approximation as:

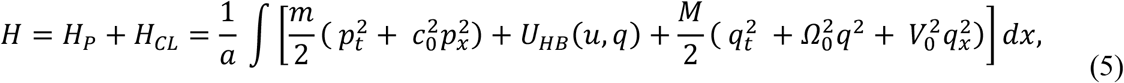

where *c*_0_ = *aω*_1_ and *V*_0_ = *l*Ω_1_.

The equations of motion for the coupled proton and CL systems with Hamiltonian *H* (5) may be written in the form:

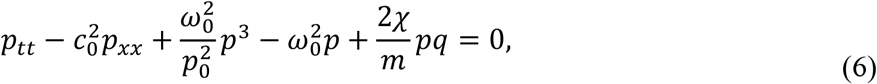

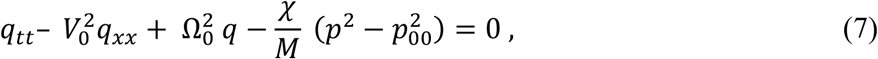

where

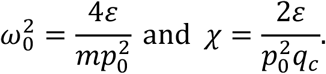

Equations of motion (6) and (7) were derived by substitution of the following derivatives of the potential *U*_*HB*_(*u*, *q*)

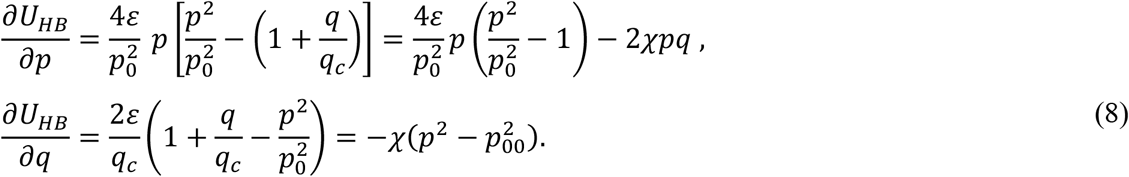

As a result, two coupled nonlinear differential equations (6) and (7) were obtained where a link between two subsystems can be derived from the Hamiltonian *H*_*int*_ of the interaction between the HB and CL chains:

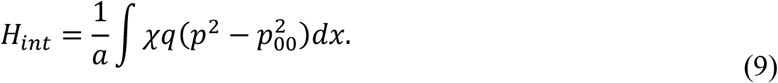

Turning to the new spatial variable ξ = *x* − *Vt* in Eqs. (6) and (7), the following set of two ordinary differential equations may be written:

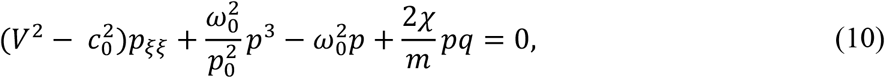

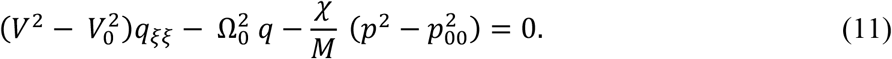

Of particular interest is the excitation moving along the chain at constant velocity *V* = *V*_0_.

Then, variable *q* can be expressed from Eq. (11):

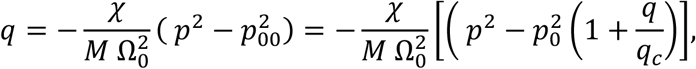

It allows expressing *q* though *p* in the form:

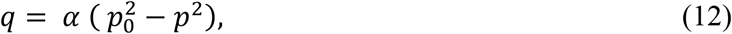

where

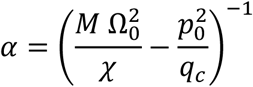

Eq. (10) after substituting Eq. (12) takes the form:

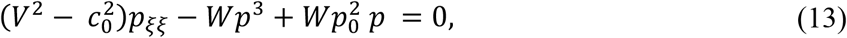

where

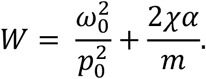

Eq. (13) can be written in a form of the well-known equation of the φ^4^-theory

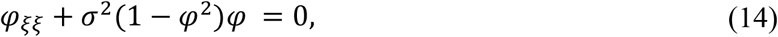

where the following variable and parameters were introduced

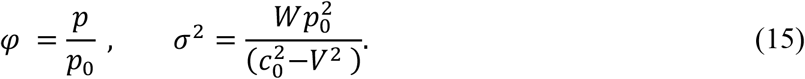

At a fixed value of velocity *V* = *V*_0_, Eq. (14) has the exact soliton solution for proton displacement in the variables *x* and *t*:

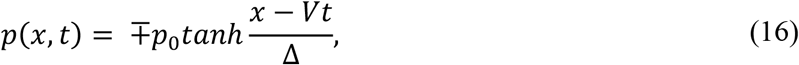

Substituting Eq. (16) into Eq. (12), we obtained a relationship for lipid headgroup displacement in the form of soliton solution:

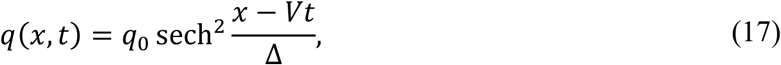

where *q*_0_ = *αp*_0_ and soliton half-width Δ is defined as an inverse value of parameter *σ*^2^:

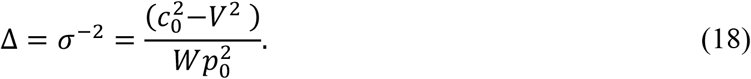

The solution *p*(*x*, *t*) (16) for Eq. (6), shown in Fig. 4A, corresponds to the soliton solution of the well-known equation of the *φ*^4^-theory (see, e.g. (Davydov 1985)) and defines a kink and antikink (upper and lower signs in Eq. (16), correspondingly). The solution *p*(*x*, *t*) defines the motion of ionic defect (hydronium) and corresponds to domain wall motion, when all protons on the left side of the domain wall are in the right-hand well of the double-well potential (3), and all protons on the right side are in the left-hand well (bottom scheme in Fig. 2). The proton (hydronium ion) transport along the HB chain corresponds to the motion of this domain wall of width Δ (16), when proton moves from the right-hand well to the left-hand well of the double-well potential (Fig. 3).

**Fig. 4.**
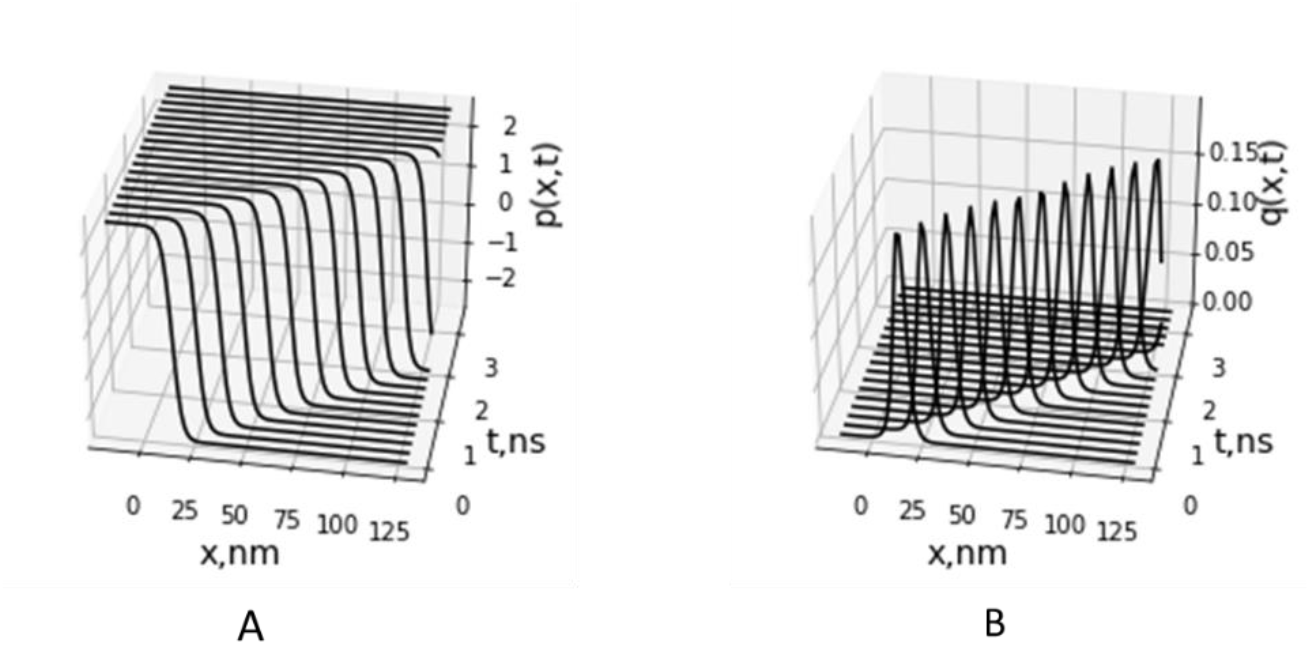
Propagation of the kink corresponding to proton displacements in the HB chain *p*(*x*, *t*) (A) and motion of soliton of deformation of CL headgroup chain *q*(*x*, *t*) (B).

The solution *q*(*x*, *t*) (17) represents the acoustic soliton (Fig. 4B) and describes the local deformation (compression) of the membrane region moving with velocity *V* in the CL chain accompanied with proton motion in the HB chain. We suggested that local compression of the lipid headgroups causes compression in the HB chain hydrogen bonding to the phosphate oxygens of the CL headgroups. This leads to a decrease in the height of the HB potential barrier due to reduction of the distance between OH^−^ groups less than critical value *R_c_*. According to Eq. (2) and Fig. 3, the HB potential at the distance *R* in the range of *R_c_* is locally transformed from double well potential to the low-barrier HB (LBHB) potential and then to the single well potential that facilitates proton transfer between donor and acceptor OH^−^ groups.

In the model, we considered a special case of soliton motion with the fixed velocity *V* = *V_0_* which is the specific velocity defined by the dispersion of lipid molecule vibration in the MIM (Eq. (4)). Soliton motion with other velocities needs further investigation of the model and computational solution of the nonlinear Eqs. (10) and (11). For similar two-component models allowing soliton solutions (Antonchenko et al. 1983; Lupichev et al. 2015), dissipationless soliton has a admissible velocity range *v* < *γc*_0_, where parameter γ <1 is defined by internal parameters of the systems. In our case, we estimated maximum soliton velocity as *V0* =50 m/s - 100 m/s using relationship *V*_0_ = *l*Ω_1_, where the estimated distance between lipid molecules on the MIM surface is *l* = 0.5 nm for surface area of CL molecules of 0.85 nm^2^ (Lemmin et al. 2013) and estimation of the characteristic frequency of the headgroup dipole oscillations being on the order of Ω1 =10^10^Hz - 10^11^ Hz. The estimated value of *V_0_* lays in the range of sound velocities 50-300 m/s measured in the lipid monolayers (Griesbauer et al. 2009) and velocity of the solitary waves observed in lipid liposomes (Heimburg and Jackson 2005). The soliton half-width value Δ ≌ 8.0 nm was estimated in a dynamics model of soliton formation and propagation in domain structures in lipid membranes (Kadantsev and Goltsov 2022). At the estimated value of width Δ, the soliton region covers approximately 10 lipid molecules. These values of soliton velocity *V_0_* =50 m/s and width were used in calculation of soliton propagation shown in Fig. 4.

## 3. Discussion

In our model, we considered the complex cristae ultra-structure and used new data on the macromolecular organization of the OXPHOS system obtained in our laboratory by cryo-electron tomography (Nesterov et al. 2021b). The ordered clustered oligomeric structure of parallel rows of respirasomes and ATP synthase dimers was observed to be located on the cristae folds of the MIM. The short distance between the observed rows of proton pumps and ATP synthase dimers provides the conditions for direct and fast transfer of protons to ATP synthases along cristae membrane. Apart from the charge transport, pumped protons can also possess Gibbs energy excess due to their interactions with lipid and water molecules on the membrane-water interface. It is expected that this energy should not be wasted during efficient function of mitochondria, but could be consumed by ATP synthase. The interest to the kinetic coupling in mitochondria (Toth et al. 2020), which is most likely provided by fast lateral proton transport, is increasing, but the molecular mechanism of this process remains to be fully understood.

To describe lateral transport of energized protons from proton pumps (sources) to ATP synthases (sinks), we developed a molecular model of long-range proton transport along the cristae membrane, which is accompanied by excitation of elastic waves moving along the membrane and facilitating proton transfer. We considered Grotthuss-like mechanism of proton motion in the HB chains formed by water molecules strongly hydrogen bonding with the phosphate oxygens at the water-lipid interface. The HB chain was suggested to be assembled on the surface of CL domains, which were observed in CL-enriched cristae membrane (Mileykovskaya and Dowhan 2009). The developed two-component dynamics model of the proton transport combined two subsystems including the HB chain of water molecules and CL polar headgroup chain. We analytically obtained two-component soliton solution in the model, which describe the motion of the proton kink, corresponding to successive proton hops from one quasi equilibrium position to another in the HB chain. The proton kink motion in HB chain is accompanied by the motion of compression soliton in the chain of CL polar headgroups. The local deformation of the membrane during the passage of the soliton facilitates proton hops due to water molecules approaching each other in the HB chain.

Experimental observation of soliton-type excitations in lipid mono- and bilayers have been carried out in a number of experiments using different methods for excitation and registration of elastic pulses. In the experiments with optical generation of solitary waves, excitation of acoustic soliton-like pulses and their dissipationless propagation have been observed in lipid monolayers at surface pressure above a certain threshold value (Shrivastava and Schneider 2014). Soliton-type excitation of elastic pulse has also been observed in liposomes in temperature region of lipid phase transition and its estimated velocity was about 115 m/s (Heimburg and Jackson 2005). When the soliton moves, the distance between the acid headgroups decreases, and compression of the lipid is up to 20%. It was proposed that nonlinearity of the membrane compressibility function (depends on temperature and pressure) in the vicinity of the melting transition provides the condition for soliton propagation (Heimburg and Jackson 2005). The sharp decrease of proton lateral diffusion coefficient after lipid phase transition into the liquid-ordered state was also reported earlier at the monolayer and was associated with hydrogen-bond network breakup (Prats et al. 1987). Interestingly, we also registered earlier that effectiveness of mitochondrial ATP synthesis increases near the temperature of lipid phase transition into liquid-disordered phase (Nesterov et al. 2014; Yaguzhinsky et al. 2017). Note, that not only the temperature and pressure can affect the phase state of the lipid membrane – it is also controlled by pH (by protons). In accordance to this, propagation of solitary waves was observed on a lipid interface at pulses generation by local acidification (Fichtl et al. 2016). The pH-induced perturbation of a lipid interface propagated at the velocities up to 1.4 m/s. The obtained correlation between the mechanical (monolayer compressibility), thermodynamic characteristics of the interface and the pulse velocities confirmed the acoustic mechanism of these pulses and proton transfer along the lipid monolayer. Fichtl et al. proposed that proton soliton-like transport offers effective enzyme-enzyme communication in cells. Note the defined velocity of solitary wave is much slower than the value of sound velocity ranging from 50 m/s to 300 m/s measured in the lipid monolayers (Griesbauer et al. 2009). Our estimete of soliton velocity of 100 m/s gave an upper limit of the velocity and the developed model suggested that proton soliton can possess a spectrum of the velocities limited by sound velocity in the lipid membranes. The further development and analysis of the model would show what velocity is achieved in the MIM.

The results of our modelling revealed the significant role of membrane deformation in the mechanism of lateral proton transfer. This finding agrees with experimental data which showed dependence of the proton transfer efficiency on the surface area of lipid headgroups and surface pressure applied to the membrane (Prats et al. 1987). The fast proton transport was reported to occur in a defined region of surface pressure and was abolished beyond its boundaries. These experimental data also confirmed our model assumption on the tight coupling between the HB networks formed by the lipid headgroups and interface water molecules. In the model, strain in the water HB chain (Eq. 6) is defined by the local deformation in the CL chains (Eq. 7) that causes formation of the two-component soliton.

In our model, we considered a significant role of CL molecules in the assembling of proton-conducting structures in the MIM. The experimental and computational works proposed that CL molecules (being unique anionic lipid), besides their essential contribution to mitochondrial structure, functions, and stress response (see section 2.2), could also play a central role in proton traps at the lipid-water interface and contribute to rapid lateral proton conduction along the membranes (Haines and Dencher 2002; Dahlberg et al. 2010). Proton-conducting structures in the MIM proposed in our model could be formed as a result of CL domain organization in the membranes which was obtained experimentally (Mileykovskaya and Dowhan 2009; Pennington et al. 2017) as well as in computational simulation (Lemmin et al. 2013; Wilson et al. 2019).

The developed model of the proton-carrying soliton described a stage of soliton propagation over membrane-water interface and did not consider the initial stage of the soliton formation. This stage should be investigated thoroughly by modelling the local perturbation of the mitochondria membrane by proton or hydroxonium. At the current step of the model development, we proposed that proton-carrying soliton might be initially excited by protons pumped to the outer side of the MIM. The local decrease of pH and incompletely compensated highly localized proton/hydroxonium charge cause weakening of electrostatic repulsion interactions between lipid headgroups that lead to an increase of local lipid packing, accompanied by a decrease in lipid surface area and local deformation of the membrane (Tokutomi et al. 1980; Angelova et al. 2018; Deplazes et al. 2018). We proposed that the mechanism of proton-induced local deformation of the membranes may be modelled based on the experimental data on the formation of inward plasma membrane curvature at decreasing local extracellular pH (Ben-Dov and Korenstein 2013). Consideration of this process would allow description of the full energy of proton/hydronium interaction with the membrane, which is stored not only in electrical field, H^+^ concentration gradient and hydration shell of the proton, but also in elastic deformation of the MIM.

Within the developed model, we made several assumptions and did not take into account a number of important features of the MIM and the OXPHOS system structures. First, we did not consider electrochemical transmembrane potential driving ATP synthesis. Introduction of the electric transmembrane potential is proposed to lead to additional stress of the membrane (Landau et al. 1995). Effect of electrical fields on membrane deformation was observed, in part, by the method of inner field compensation on the bilayer membranes. Its application showed a change of the membrane thickness and capacitance when the membrane interacts with various compounds (Abidor I.G. et al. 1980; Cherny et al. 1980; Jiménez-Munguía et al. 2019), including the membrane-bound proton fraction (Antonenko et al. 1993; Tashkin et al. 2019). We proposed that a similar effect may also present in the MIM under proton pump functioning (Nesterov et al. 2022). Consideration of the additional deformation of the MIM in the modified model would allow investigation of the peculiarities of the proton transport along the strained MIM.

Second, a significant factor defining lateral proton motion along the MIM is the local pH gradient of 0.3 units experimentally observed between proton pumps and ATP synthases (Rieger et al. 2014). Introduction of the corresponding electric field as lateral proton motive force into the model (in right hand side of Eq. (6)) would lead to unidirectional motion of proton-carrying soliton from the proton pumps to ATP synthases.

Third, the developed model did not take into account restoration of water molecule orientation in the HB chain after proton passage. This did not allow describing continuous proton transfer in the HB chain at the water-membrane interface. As seen in the bottom scheme in Fig. 2, proton transfer along the HB chain reorients the chain configuration, which should be restored in order to reset transfer of the next proton. The restoration of the initial orientation of the HB chain significantly hinders proton transport following Grotthus-like mechanism. As reported, fast proton hops between sites (1 ps) according to Grotthus-like mechanism is followed by much slower reorientation of the HB chain (100 ps) (Pomès and Roux 1998; Wraight 2006). To overcome this problem, additional mechanisms were suggested to reset the HB chain in the soliton models of proton transport and describe continuous proton transfer in the HB chain (Pnevmatikos 1988). We proposed that the energy of water molecule reorientation might be provided by electrochemical potential across the MIM as well as pH gradient along the MIM. These factors may accelerate relaxation of the HB chain to preferable orientation of electric dipoles of water molecules in the lateral electric field induced by the pH gradient along MIM (Fig. 2A). Consideration of these factors in the model at the further stage of the model development will allow description of the continuous proton conductivity through the HB chains of water molecules between proton pumps and ATP synthases along the MIM.

## Conclusion

Based on the experimental data on cryo-electron tomography and nonlinear dynamics modelling, we developed the model-based approach to the investigation of the coupled proton and energy transport along mitochondria crista membranes and proposed that proton conductive networks provide a direct connection between rows of proton pumps and ATP synthases assembled on the crista folds. The developed model describes collective excitation as two-component soliton which can be formed and propagate in the strongly linked systems of the hydrogen bond chains of ordered water molecules and membrane lipid polar headgroups. We suggested that the advantage of the proposed soliton-type mechanism lies in achieving coherent transport of proton and energy, which can be simultaneously utilized by ATP synthase. As we established in the model, the crista membrane, its structure, topology, and lipid composition play a significant role in proton and energy transport and communication between key components of the OXPHOS systems. The further investigation of the molecular mechanism of the direct proton and energy transport in mitochondria crista needs detailed consideration of the proton environment including a unique system of the water-membrane interface in the OXPHOS system. Experimental verification of the proposed soliton-type communication mechanism in mitochondria could be carried out by investigation of the effect of proton environment modulation on the effectiveness of ATP synthesis. Another way to verify the proposed mechanism of the coupled proton and energy transport along mitochondria membrane is the use of bio-inspired biotechnology in the investigation of ATP synthesis effectiveness in giant liposomes or in other artificial systems encapsulating ATP synthases and mimicking mitochondrial environment.

## Author Contributions

Conceptualization, L.S. Y., G.W. and V.N.K.; methodology, S.V.N. and A.N.G.; writing—original draft preparation, S.V.N. and A.N.G.; writing—review and editing, L.S. Y., G.W., V.N.K., R.G.V., S.V.N. and A.N.G.; supervision, R.G.V. All authors have read and agreed to the submitted version of the manuscript. S.V.N. and A.N.G. contributed equally to this work.

